# A defect in thymic tolerance causes T cell-mediated autoimmunity in a murine model of COPA syndrome

**DOI:** 10.1101/2020.01.21.914440

**Authors:** Zimu Deng, Christopher S. Law, Frances O. Ho, Kristin M. Wang, Kirk D. Jones, Jeoung-Sook Shin, Anthony K. Shum

## Abstract

COPA syndrome is a recently described Mendelian autoimmune disorder caused by missense mutations in the *Coatomer protein complex subunit alpha (COPA)* gene. Patients with COPA syndrome develop arthritis and lung disease that presents as pulmonary hemorrhage or interstitial lung disease (ILD). Immunosuppressive medications can stabilize the disease, but many patients develop progressive pulmonary fibrosis that requires life-saving measures such as lung transplantation. Because very little is understood about the pathogenesis of COPA syndrome, it has been difficult to devise effective treatments for patients. To date, it remains unknown which cell types are critical for mediating the disease as well as the mechanisms that lead to autoimmunity. To explore these issues, we generated a *Copa^E241K/+^* germline knock-in mouse bearing one of the same *Copa* missense mutations in patients. Mutant mice spontaneously developed ILD that mirrors lung pathology in patients, as well as elevations of activated cytokine-secreting T cells. Here we show that mutant Copa in epithelial cells of the thymus impairs the thymic selection of T cells and results in both an increase in autoreactive T cells and decrease in regulatory T cells in peripheral tissues. We demonstrate that T cells from *Copa^E241K/+^* mice are pathogenic and cause ILD through adoptive transfer experiments. In conclusion, we establish a new mouse model of COPA syndrome to identify a previously unknown function for Copa in thymocyte selection and demonstrate that a defect in central tolerance is a putative mechanism by which *COPA* mutations lead to autoimmunity in patients.

**One Sentence Summary:** A new mouse model of COPA syndrome develops lung pathology that recapitulates patients and reveals that T cells are important drivers of the disease.

## Introduction

Our lab recently co-discovered COPA syndrome, a Mendelian autoimmune disorder of inflammatory arthritis and interstitial lung disease (ILD) caused by dominant mutations in the *Coatomer subunit alpha* (*COPA*) gene (*1*). COPA syndrome typically presents in childhood with arthritis or pulmonary disease that manifests as ILD or life-threatening pulmonary hemorrhage. Although most patients usually receive long-term immunosuppression, all eventually develop ILD that typically begins as follicular bronchiolitis and over time evolves to include cystic disease. Many if not all patients experience a progressive decline in lung function that may eventually advance to end-stage pulmonary fibrosis forcing some patients to undergo lung transplantation (*2*). Because COPA syndrome was only recently described, the most effective therapies for combating the disorder have not been defined. Moreover, it remains unclear whether early recognition and treatment can prevent some of the deadlier complications of the disease.

One of the major challenges to devising therapies tailored to COPA syndrome has been that very little is understood about the pathogenesis of disease. COPA is a subunit of coat protein complex I (COPI) that mediates retrograde movement of proteins from the Golgi apparatus to the endoplasmic reticulum (*3*). Because COPA is ubiquitously expressed, it is not known which cell types are important for initiating or propagating the disease (*2, 4*). Furthermore, the mechanisms by which mutations in *COPA* lead to a breakdown in immunological tolerance and autoimmunity remain a mystery. To explore these questions, we generated a *Copa^E241K/+^* germline knock-in mouse bearing one of the same *COPA* missense mutations in patients. Mutant mice spontaneously developed lung disease and systemic inflammation mirroring some of the key clinical features of patients (*1*). Here, we used *Copa^E241K/+^* mice to define the mechanisms that trigger autoimmunity and to identify critical cell types responsible for mediating disease. Through our work, we present a novel translational model to further investigate the molecular mechanisms of COPA syndrome and develop treatments for patients suffering from this highly morbid disease.

## Results

### Copa^E241K/+^ mice recapitulate lung disease in patients

Prior work identified four missense mutations in COPA syndrome each of which result in a unique amino acid change in COPA that is 100% conserved to yeast. We generated a germline point mutant knock-in mouse to express the E241K amino acid change found in one COPA syndrome family (*1*) (fig. S1A-F). Resulting mice crossed to C57Bl/6J wild-type (WT) mice were grossly normal and fertile. Breeding of heterozygous *Copa^E241K/+^* mice did not result in homozygous *Copa^E241K/E241K^* mice. To determine whether *Copa^E241K/+^* mice developed clinical features of COPA syndrome patients, we analyzed aged animals for disease. None of the mice developed appreciable erythema or swelling of the ankle, foot or digits consistent with inflammatory arthritis. Histological examination confirmed the absence of immune cell infiltration or cartilage loss in the joints (fig. S2A-B). Lungs harvested from aged mice, however, revealed spontaneous disease that nearly matched the lung pathology described in COPA syndrome patients (Fig. 1A-N and fig. S2C-D) (*1, 2*). In general, *Copa^E241K/+^* mice exhibited cellular bronchiolitis with germinal center formation, the most common pathologic finding in lung biopsies from COPA syndrome patients (Fig. 1D-E). Immunohistochemical staining revealed mononuclear cell infiltrates containing a predominance of CD4^+^ T cells and lymphoid aggregates comprised of B cells (fig. S2D). The immune infiltrates were in a pattern nearly identical to similar staining performed in patients (*1*).

**Figure 1.**
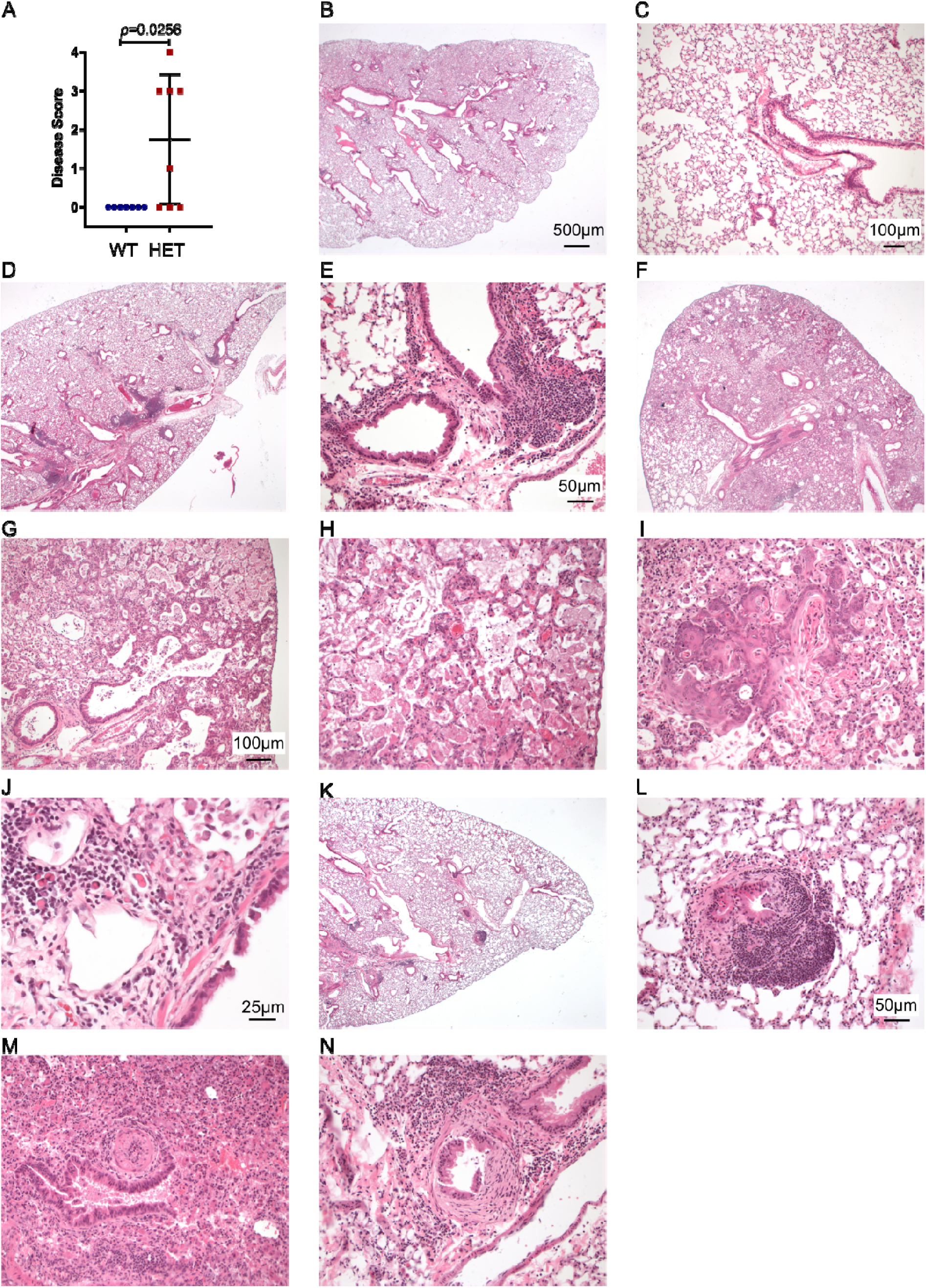
*Copa^E241K/+^* mice spontaneously develop lung disease. Shown are the mouse lung disease scores and select images from 1 *Copa^+/+^* (WT, panels B-C) and 3 *Copa^E241K/+^* (HET-1, panels D-E; HET-2 panels F-J; HET-3 panels K-N) mice. A. Disease scores of lung sections from WT and HET mice. (10-11-month-old littermates: WT, *n* = 7; HET, *n* = 8). B. Low power image of a WT showing normal pulmonary architecture without significant inflammation, consolidation, or fibrosis. C. Higher power image of the WT lung. D. Low power image of HET-1 showing lymphocytic peribronchial and peribronchiolar inflammation. E. Higher power image of HET-1 showing peribronchiolar inflammation. F. Low power image of HET-2 showing prominent consolidation of distal alveoli by edema, fibrin, proteinosis, and macrophage accumulation. G. A higher power image of HET-2 showing alveolar septal thickening with edema, type 2 pneumocyte hyperplasia, and mild mixed acute and chronic inflammation. H. A high power image of HET-2 showing alveolar spaces filled by macrophages with foamy cytoplasm and proteinaceous granular fluid. I. Another high-power image of HET-2 showing the terminal bronchioles and alveolar ducts with prominent squamous metaplasia and keratinized epithelium being shed into air spaces. J. A 40x view of HET-2 showing the peribronchiolar inflammation with a mixed infiltrate of lymphocytes and plasma cells, including Mott cells (plasma cells with cytoplasmic immunoglobulin inclusions). K. On low power of HET-3, patchy peribronchial lymphoid hyperplasia is noted. L. Higher power image of HET-3 showing a lymphoid nodule in the bronchiolar subepithelium. Focal epithelial denudation is present with granulation tissue extending into the lumen (follicular bronchiolitis with proliferative bronchiolitis). M. Higher power image of HET-3 showing a pulmonary artery with marked luminal narrowing by endothelial hyperplasia and mild chronic inflammation. N. Higher power image of another bronchiole of HET-3 showing prominent eccentric proliferation of fibroblasts in the subepithelium (but above the muscular layer), diagnostic of constrictive bronchiolitis. Mann–Whitney U test was used in A. p < 0.05 is considered statistically significant. ns: not significant. Images B, D, F, K taken at 2x magnification, scale bar = 500 μM; images C, G, H, I taken at 10x, scale bar = 100 μM. Images E, L, M, N taken at 20x, scale bar = 50 μM. Image J taken at 40x, scale bar = 25 μM.

Further analysis of lung pathology revealed a spectrum of acute and chronic lung damage that ranged from mild to severe (Fig. 1D-N). One mouse had evidence of acute lung injury and diffuse alveolar damage demonstrated by distal alveolar consolidation with edema, fibrin, proteinosis and macrophage accumulation (Fig. 1F-J). Higher power views showed type 2 pneumocyte hyperplasia (Fig. 1G) and peribronchiolar inflammation with mixed infiltrates of lymphocytes that included Mott cells (B cells with cytoplasmic inclusions of immunoglobulins) that have been described in other autoimmune mouse models (Fig. 1J) (*5*). Although we did not observe frank pulmonary hemorrhage, one mouse had mild inflammation in a pulmonary artery suggesting vasculitis (Fig. 1M) as well as evidence of lung fibrosis in an area of constrictive bronchiolitis (Fig. 1N). Thus, despite the absence of joint inflammation, *Copa^E241K/+^* mice spontaneously developed ILD, the organ manifestation that appears to most strongly impact the prognosis and clinical course of COPA syndrome patients (*2*).

### Copa^E241K/+^ mice spontaneously develop activated, cytokine-secreting T cells

To determine if the immune system in *Copa^E241K/+^* mice paralleled patients, we sacrificed animals at 3 months. Spleens from WT and *Copa^E241K/+^* mice demonstrated no differences in total splenocytes including absolute numbers and percentages of B and T cells (fig. S3A-C). B cells in *Copa^E241K/+^* mice had significantly elevated levels of CD86 (fig. S3D) although serum IgG levels were not increased (fig. S3E). We were unable to detect an increase in antinuclear autoantibodies (ANA) including autoantibodies to double stranded DNA or ribonuclear proteins (fig. S3F-G).

Despite the lack of circulating autoantibodies, immunophenotyping of T cells revealed several abnormalities. We observed a significant decrease in the percentage of CD44^lo^CD62L^hi^ naïve CD4^+^ and CD8^+^ T cells in *Copa^E241K/+^* mice and a significant increase in the percentages of activated CD44^hi^CD62L^lo^ effector memory (EM) CD4^+^ and CD8^+^ T cells (Fig. 2A-B). In addition, peripheral CD4^+^ and CD8^+^ T cells in *Copa^E241K/+^* mice stimulated with phorbol 12-myristate 13-acetate (PMA) and ionomycin demonstrated a significant increase in IFNγ and IL-17A-secreting CD4^+^ T cells and IFNγ-secreting CD8^+^ T cells (Fig. 2C-D). Thus, introducing the E241K *Copa* point mutation into mice triggered early, spontaneous inflammation with activation of cytokine-secreting CD4^+^ T cells and interstitial lung disease that phenocopied similar clinical features in COPA syndrome patients (*1*).

**Figure 2.**
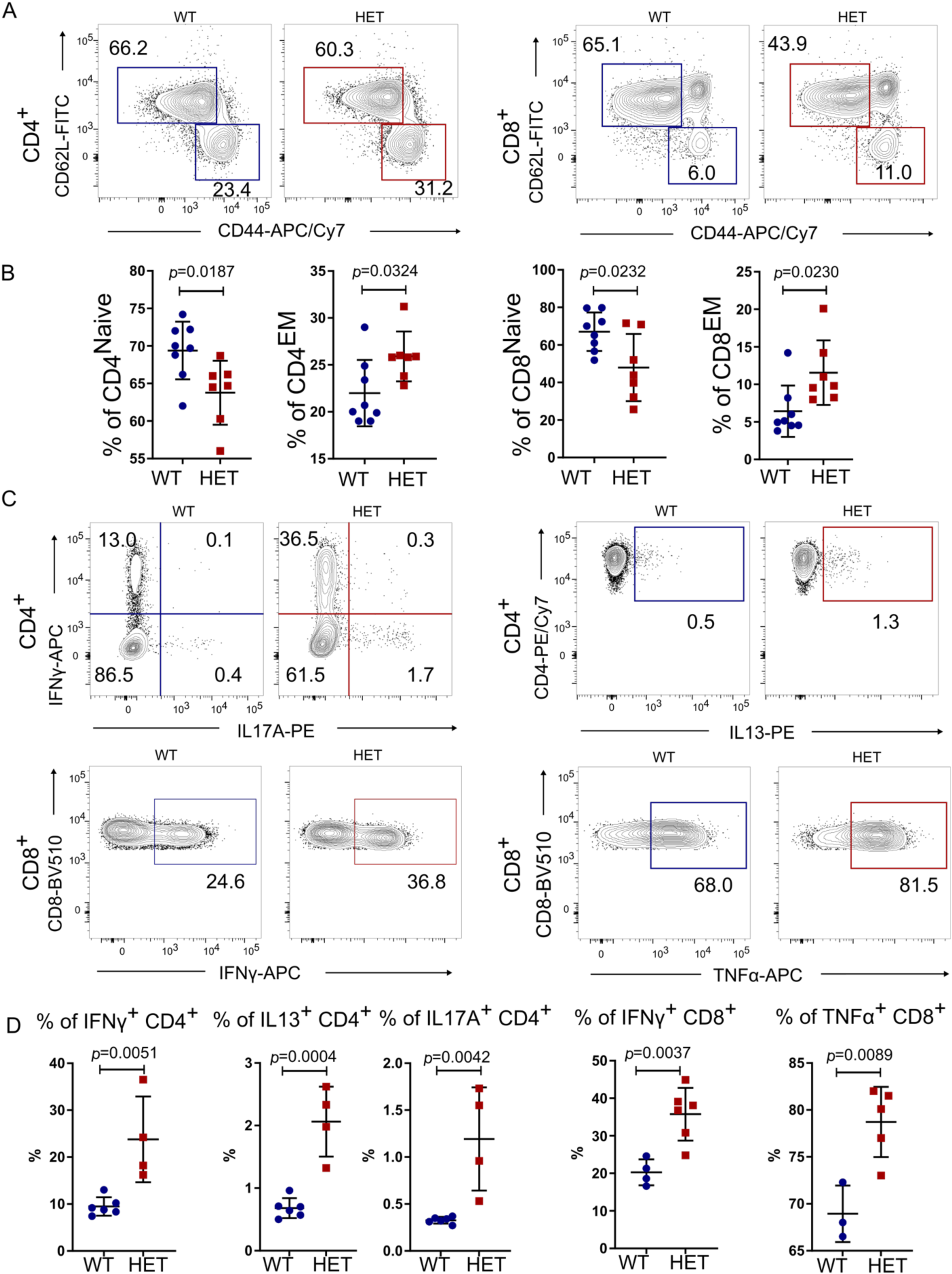
*Copa^E241K/+^* mice have spontaneous activation of cytokine-secreting T cells. A. Representative flow plots of CD62L versus CD44 expression on splenic CD4^+^ and CD8^+^ T cells from *Copa^+/+^* (WT) and *Copa^E241K/+^* (HET) mice. B. Percentages of naïve and memory CD4^+^ (left) and CD8^+^ T (right) cells from indicated mice (3-month-old littermates: WT, *n* = 8; HET, *n* = 7). C. Representative flow plots of intracellular cytokine levels in splenic CD4^+^ (top: IFNγ, IL17A and IL13) and CD8^+^ T cells (bottom: IFNγ and TNFα) after PMA/Ionomycin stimulation. D. left: Percentages of IFNγ, IL17A and IL13 producing CD4^+^ T cells among total CD4^+^ T cells from indicated mice (3-month-old littermates: WT, *n* = 6; HET, *n* = 4). right: Percentages of IFNγ and TNFα-producing CD8^+^ T cells among total CD8^+^ T cells (3-month-old littermates: WT, n = 4; HET, n = 6) Data are mean ± SD. Unpaired, parametric, two-tailed Student’s t-test was used for statistical analysis. *p* < 0.05 is considered statistically significant. ns: not significant.

### Single-positive thymocytes are significantly increased in Copa^E241K/+^ mice

Given the normal B cell numbers in *Copa^E241K/+^* mice and absence of autoantibodies, we turned our focus to T cells given their central role in orchestrating autoimmunity. To determine whether the spontaneous expansion of activated peripheral T cells in *Copa^E241K/+^* mice might be related to changes in T cell development or selection we performed a detailed analysis of thymocyte populations. Total cell numbers in thymi from 4-6 weeks old *Copa^E241K/+^* and WT littermates were similar and thymocyte plots demonstrated equivalent percentages and absolute cell numbers of CD4^-^CD8^-^ double negative (DN) cells (Fig. 3A-B). Interestingly, the percentage of CD4^+^CD8^+^ double positive (DP) cells were significantly decreased and the percentages of CD4^+^ single-positive (CD4SP) and CD8^+^ SP (CD8SP) cells were significantly increased in *Copa^E241K/+^* mice (Fig. 3A). Analysis of cell counts at each thymocyte stage revealed the increased ratio of SP cells in *Copa^E241K/+^* mice was caused by an expansion of the SP cell population because DN and DP cell numbers were equivalent in *Copa^E241K/+^* and WT littermates (Fig. 3B). To identify at which stage of thymocyte selection the increase in SP cells occurred, we evaluated thymocytes by TCRβ and CD69 and found a significant increase in the percentages of TCRβ^+^CD69^+^ and TCRβ^+^CD69^-^ SP cells that arise post-positive selection(Fig. 3C) (*7, 8*). Taken together, the increase in SP cells suggests mutant Copa alters T cell development or selection as a precursor to spontaneous inflammation.

**Figure 3.**
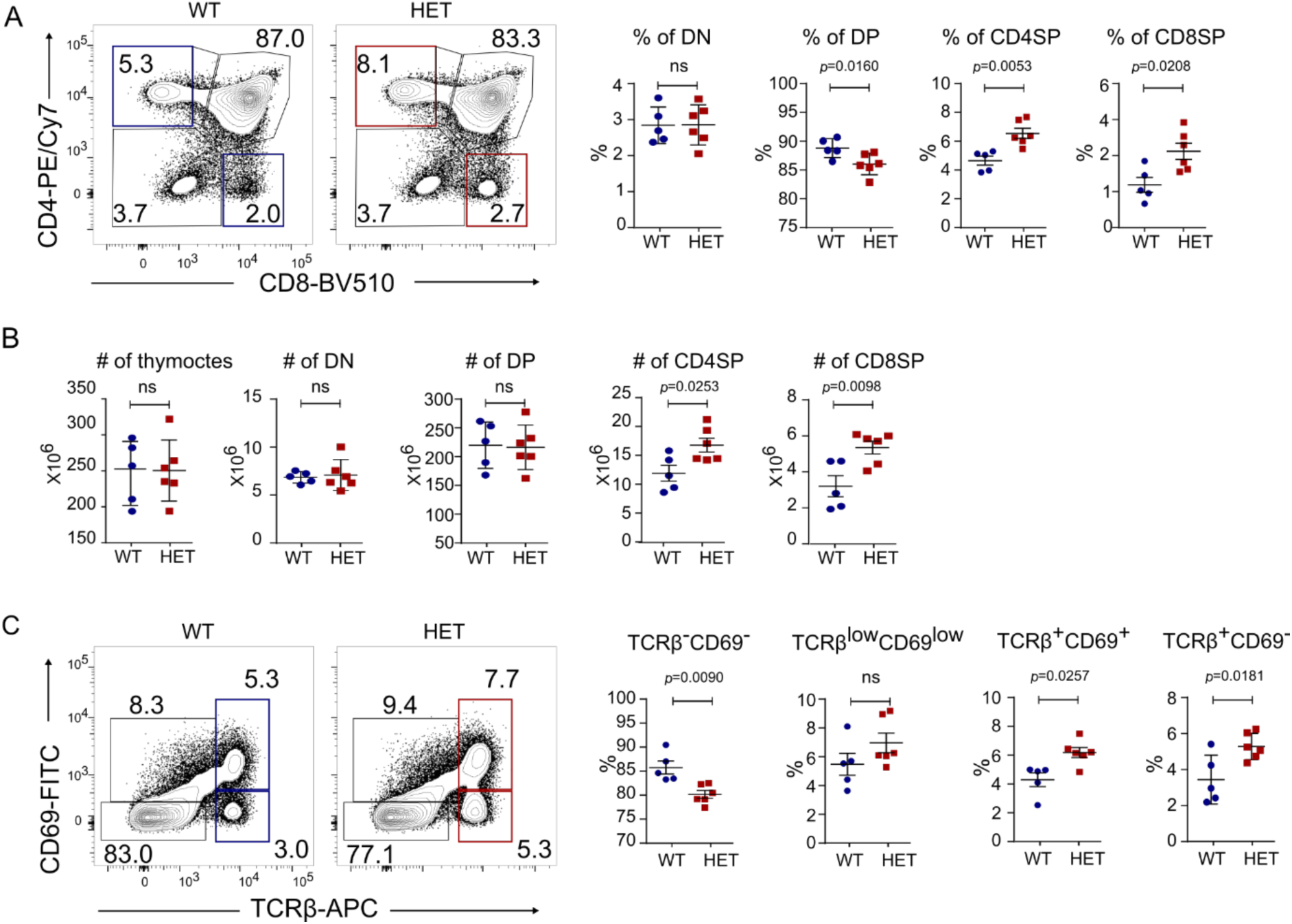
*Copa^E241K/+^* mice have increased single positive thymocytes. A. left: CD4 and CD8 profile of thymocytes. right: Percentages of double negative (DN), double positive (DP), CD4 single positive (CD4SP) and CD8 single positive (CD8SP) thymocytes from indicated mice (4-6-week-old littermates: WT, *n* = 5; HET, *n* = 6). B. Cell counts of total thymocytes, DN, DP and SP thymocytes in indicated mice (4-6-week-old littermates: WT, *n* = 5; HET, *n* = 6). C. left: CD69 and TCRβ chain on total thymocytes. right: Percentages of TCRβ^-^CD69^-^, TCRβ^low^CD69^low^, TCRβ^+^CD69^+^ and TCRβ^+^CD69^-^ populations from indicated mice (4-6-week-old littermates: WT, *n* = 5; HET, *n* = 6). Data are mean ± *SD*. Unpaired, parametric, two-tailed Student’s *t*-test was used for statistical analysis. *p* < 0.05 is considered statistically significant. ns: not significant.

### The thymic stroma causes the increased SP cells in Copa^E241K/+^ mice

Because COPA is ubiquitously expressed (*3*), we sought to understand whether the alterations in thymocyte populations was due to mutant *Copa* in thymocytes, the thymic epithelium or both. To distinguish between the contributions of hematopoietic cells and radioresistant thymic epithelial cells (TEC) we introduced congenically marked bone marrow (BM) from WT or *Copa^E241K/+^* mice into irradiated WT or *Copa^E241K/+^* hosts (*9*). Interestingly, we found an increase in CD4^+^ and CD8^+^ SP cells and mature TCRβ^+^CD69^-^ SP cells analogous to unmanipulated *Copa^E241K/+^* mice only in thymi harvested from *Copa^E241K/+^* host mice regardless of whether they had been reconstituted with WT or mutant BM (Fig. 4A-C). We also found a significant increase in effector memory T cells and lung infiltrates in *Copa^E241K/+^* host mice including those receiving WT hematopoietic cells (Fig. 5A-C), indicating that E241K Copa thymi may also contribute to the expansion of EM T cells and pulmonary disease. To evaluate definitively whether the thymic epithelium is essential to bring about the observed changes in thymocyte populations, we transplanted fetal thymic stroma harvested from WT or *Copa^E241K/+^* mice under the kidney capsule of athymic *Foxn1^nu^* nude mice (fig. S4A) (*10, 11*). Analysis of transplanted mice again revealed a significant increase in the percentages of CD4SP cells (fig. S4B) and mature TCRβ^+^CD69^-^ SP cells (fig. S4C) in *Copa^E241K/+^* thymi. Taken together, our results demonstrate that mutant *Copa* in the thymic epithelium is necessary and sufficient to cause the increase in SP thymocytes in *Copa^E241K/+^* mice and may be a critical factor in the development of lung disease.

**Figure 4.**
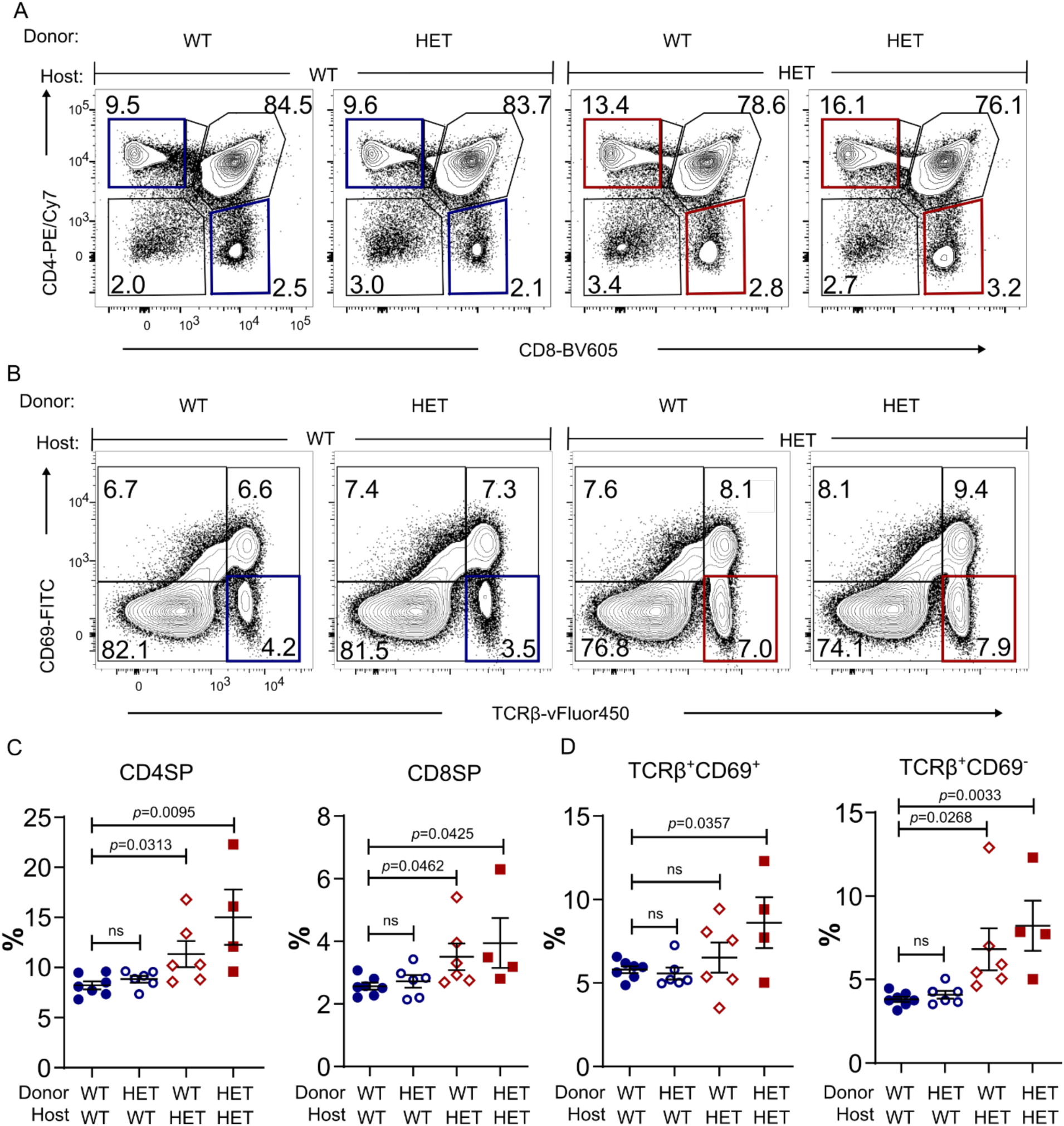
Bone marrow chimeras reveal a functional role for mutant Copa in the thymic stroma. A. Representative flow analysis of CD4 and CD8 on reconstituted thymocytes in the bone marrow chimeras. B. Flow analysis of CD69 and TCRβ expression on the reconstituted thymocytes. C. Percentages of CD4SP and CD8SP thymocytes among the reconstituted thymocytes (WT→WT, *n* = 7; WT→HET, *n* = 6; HET→WT, *n* = 6; HET→HET, *n* = 4). D. Percentages of TCRβ^+^CD69^+^ and TCRβ^+^CD69^-^ population among the reconstituted thymocytes (WT→WT, *n* = 7; WT→HET, *n* = 6; HET→WT, *n* = 6; HET→HET, *n* = 4). Data are mean ± *SD*. Unpaired, parametric, two-tailed Student’s *t*-test was used for statistical analysis. *p* < 0.05 is considered statistically significant. ns: not significant.

**Figure 5.**
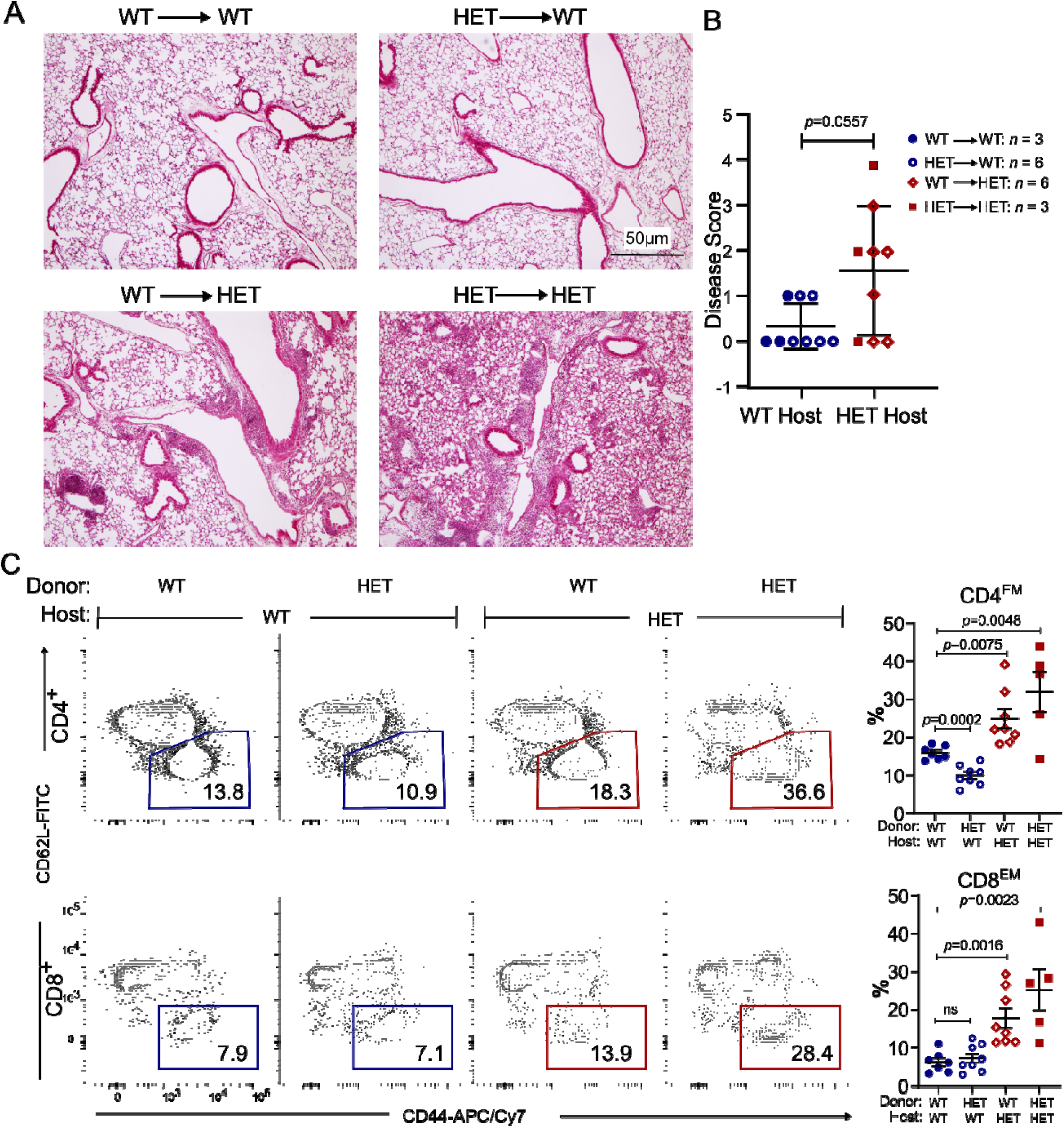
Lung infiltrates and effector memory T cells are increased in *Copa^E241K/+^* host mice. A. Representative H&E stain of the lungs from bone marrow chimeras. Scale bar: 500 μM. B. Disease scores of lung sections from bone marrow chimeras (WT→WT, *n* = 3; WT→HET, *n* = 6; HET→WT, *n* = 6; HET→HET, *n* = 3). C. left: Flow plots showing the expression of CD62L and CD44 on reconstituted splenic T cells. right: Percentages of effector memory T cell populations among the reconstituted T cells in spleen (WT→WT, *n* = 7; WT→HET, *n* = 8; HET→WT, *n* = 8; HET→HET, *n* = 5). Data are mean ± *SD*. Mann–Whitney U test was used in B. *p* < 0.05 is considered statistically significant. ns: not significant. A taken at 2x magnification, scale bar = 50µm.

### Positive selection appears unaffected in Copa^E241K/+^ *mice*

Having established that mutant COPA has a functional role in the thymic epithelium, we next sought to evaluate whether *Copa^E241K/+^* mice have a defect in the positive and negative selection of T cells. Alterations in T cell selection have been shown in multiple model systems and in human disease as an important mechanism in the pathogenesis of autoimmunity (*12*). Importantly, thymic selection may be affected not only because of a T cell-intrinsic defect, it may also occur as a result of mutations in genes that cause an abnormality within the thymic epithelium (*9, 13*).

To carefully study the impact of thymic Copa on T cell selection we employed TCR transgenic mice to track the development and selection of MHC-restricted T cells (*8, 13*). To exclude potential cell-intrinsic effects of mutant Copa on T cells, we generated chimeric mice by transplanting BM from wild-type OT-I (*14*) (H-2K^b^), OT-II (*15*) (IA^b^) and P14 (*16*) (H-2D^b^) TCR transgenic mice into irradiated *Copa^E241K/+^* and WT hosts. Analysis of thymocytes demonstrated no significant differences in the percentages of major populations between *Copa^E241K/+^* and WT hosts (Fig. 6C and fig. S4D-F). In addition, the percentages of clonotype-positive SP cells in *Copa^E241K/+^* and WT hosts were equivalent. Although these data suggest that mutant Copa in the thymic epithelium does not significantly alter the positive selection of these particular TCRs, it remains possible that Copa does have an effect on the positive selection of other TCRs within the total T cell repertoire.

**Figure 6.**
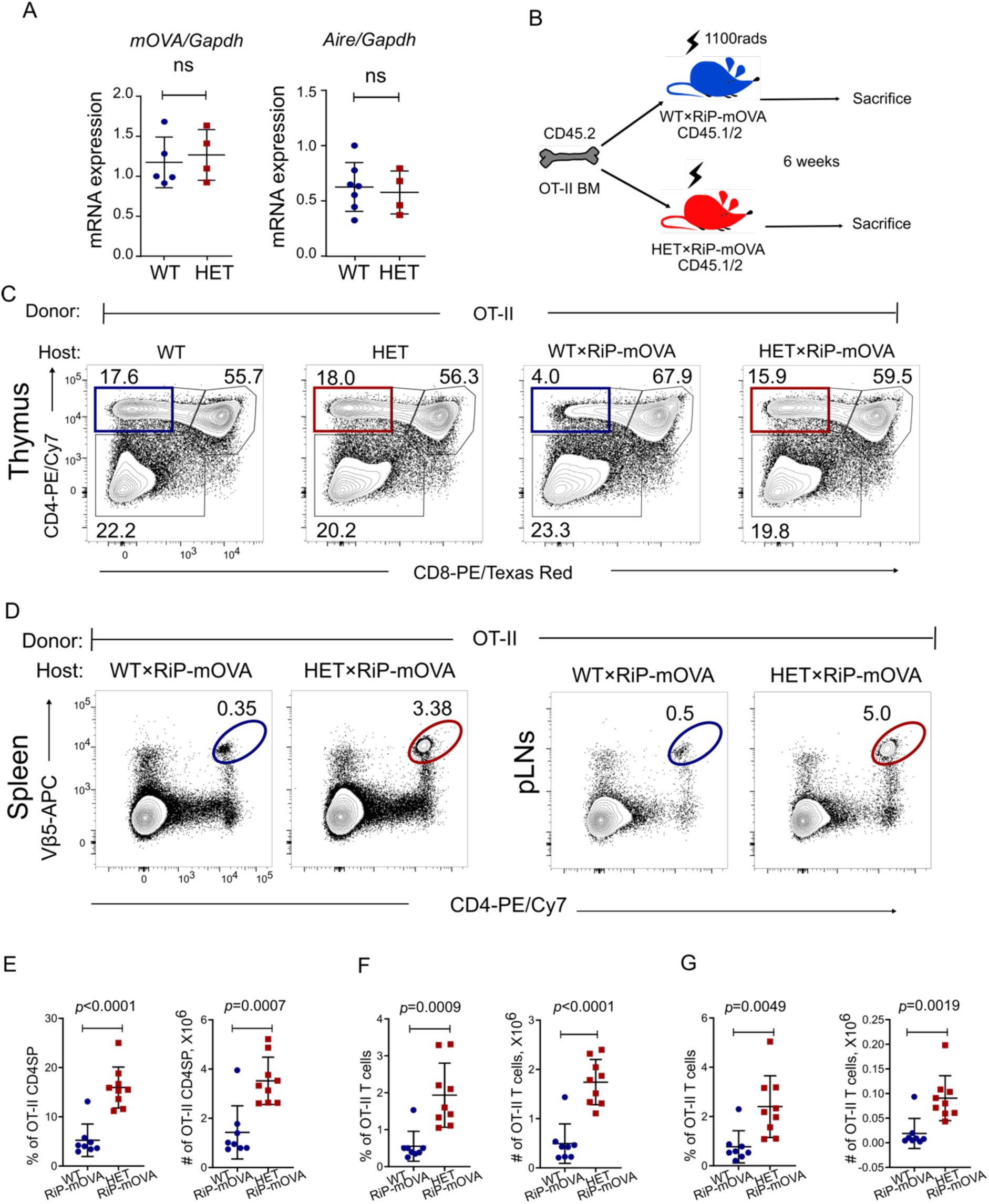
Mutant Copa impairs thymocyte negative selection and allows autoreactive T cells to escape to the periphery. A. left: *mOVA* expression level in thymic epithelial cells (4-5-week-old littermates: WT×RiP-mOVA, *n* = 5; HET×RiP-mOVA, *n* = 4). right: : mRNA levels of *Aire* in thymic epithelial cells (4-5-week-old littermates: WT, *n* = 7; HET, *n* = 4). B. Schematic for bone marrow chimeras. C. Representative flow analysis of CD4 and CD8 profile of OT-II thymocytes in chimeric mice. D. Representative flow plots showing the presence of OT-II CD4^+^ T cells in spleen (left) and pancreatic lymph nodes (right). E. Percentages (left) and absolute numbers (right) of CD4SP in the reconstituted thymi (WT×RiP-mOVA, *n* = 8; HET×RiP-mOVA, *n* = 9). F. Percentage and absolute numbers of OT-II CD4^+^ T cells in the reconstituted spleen (WT×RiP-mOVA, *n* = 8; HET×RiP-mOVA, *n* = 9). G. Percentage and absolute numbers of OT-II CD4^+^ T cells in reconstituted pancreatic lymph nodes (WT×RiP-mOVA, *n* = 8; HET×RiP-mOVA, *n* = 9). Data are mean ± *SD*. Unpaired, parametric, two-tailed Student’s *t*-test was used for statistical analysis. *p* < 0.05 is considered statistically significant. ns: not significant.

### A defect in negative selection leads to escaped autoreactive T cells in Copa^E241K^ mice

After observing no significant alteration in positive selection in *Copa^E241K/+^* mice, we next sought to assess whether they exhibited a defect in negative selection that may contribute to the increase in SP thymocytes. Importantly, a defect in thymocyte negative selection would establish a mechanism for autoimmunity in our model (*12*). To study this, we used RiP-mOVA transgenic mice bred to OT-II TCR transgenic mice, a standard approach to studying the selection of T cells to a thymically-expressed self-antigen (*17*). RiP-mOVA mice express membrane bound ovalbumin (mOVA) as a neo-self-antigen specifically within medullary thymic epithelial cells (mTECs). OT-II cells express a TCR specific for ovalbumin peptide. Under normal conditions, OT-II cells undergo negative selection and die upon encountering ova in the thymus of RiP-mOVA mice.

First, we harvested thymic epithelial cells from double transgenic mice to assess whether mutant Copa altered mRNA expression of the mOVA neo-self-antigen. Quantitative PCR showed that *mOVA* mRNA levels were equivalent between WT and mutant mice (Fig. 6A). Similarly, we confirmed there was no difference in expression of the transcriptional regulator Aire (Fig. 6A), since loss of *Aire* has previously been shown to impact selection of T cells to mOVA(*17, 18*). Having confirmed that mutant Copa did not affect *mOVA* or *Aire* expression, we next transplanted BM from WT OT-II mice into RiP-mOVA*/Copa^+/+^* and RiP-mOVA/*Copa^E241K/+^* mice and analyzed thymocyte negative selection (Fig. 6B). We observed the expected deletion of clonotype-positive OT-II thymocytes in RiP-mOVA*/Copa^+/+^* mice due to their encounter with cognate self-antigen when compared to *Copa^+/+^* mice without the RiP-mOVA transgene (*17*) (Fig. 6C, E). In contrast, expression of E241K Copa in the thymus of RiP-mOVA/*Copa^E241K/+^* mice caused a striking impairment in negative selection that lead to a substantial increase in the absolute cell number and percentage of autoreactive, clonotype-positive SP cells (Fig. 6C, E) when compared to RiP-mOVA*/Copa^+/+^* mice. To evaluate whether self-reactive clones escaped the RiP-mOVA/*Copa^E241K/+^* thymus and entered the peripheral circulation, we harvested lymphocytes from spleen and lymph nodes of chimeric mice. Indeed, we found that RiP-mOVA/*Copa^E241K/+^* mice (Fig. 6D, F-G) had significantly increased OT-II T cells in secondary lymphoid organs in comparison to controls. Taken together, our results show that mutant Copa in the thymic epithelium causes a significant defect in thymocyte negative selection that allows for the release of autoreactive T cells into peripheral tissues capable of causing organ autoimmunity.

### Antigen-specific regulatory T cells are decreased

Having shown that *Copa^E241K/+^* mice fail to purge the immune system of autoreactive T cells, we next sought to determine the impact that mutant Copa had on Foxp3^+^CD4^+^ regulatory T cells (Tregs). We pursued these experiments given the important role that thymic selection plays in the generation of Tregs (*19*) and because Tregs are an immunosuppressive cell population critical for controlling autoimmunity (*20*). Interestingly, examination of thymi from RiP-mOVA/*Copa^E241K/+^* chimeras reconstituted with WT OT-II cells revealed that clonotype-positive Foxp3^+^CD4^+^ T cells were significantly reduced (Fig. 7A). We also found that the number of clonotype-positive Foxp3^+^CD4^+^ T cells were substantially decreased in peripheral lymphoid organs (Fig. 7B-C). Thus, in parallel to the elevations of circulating autoreactive T cells, *Copa^E241K/+^* mice showed a remarkable loss of immunosuppressive antigen-specific Tregs needed for limiting inflammation and tissue damage.

**Figure 7.**
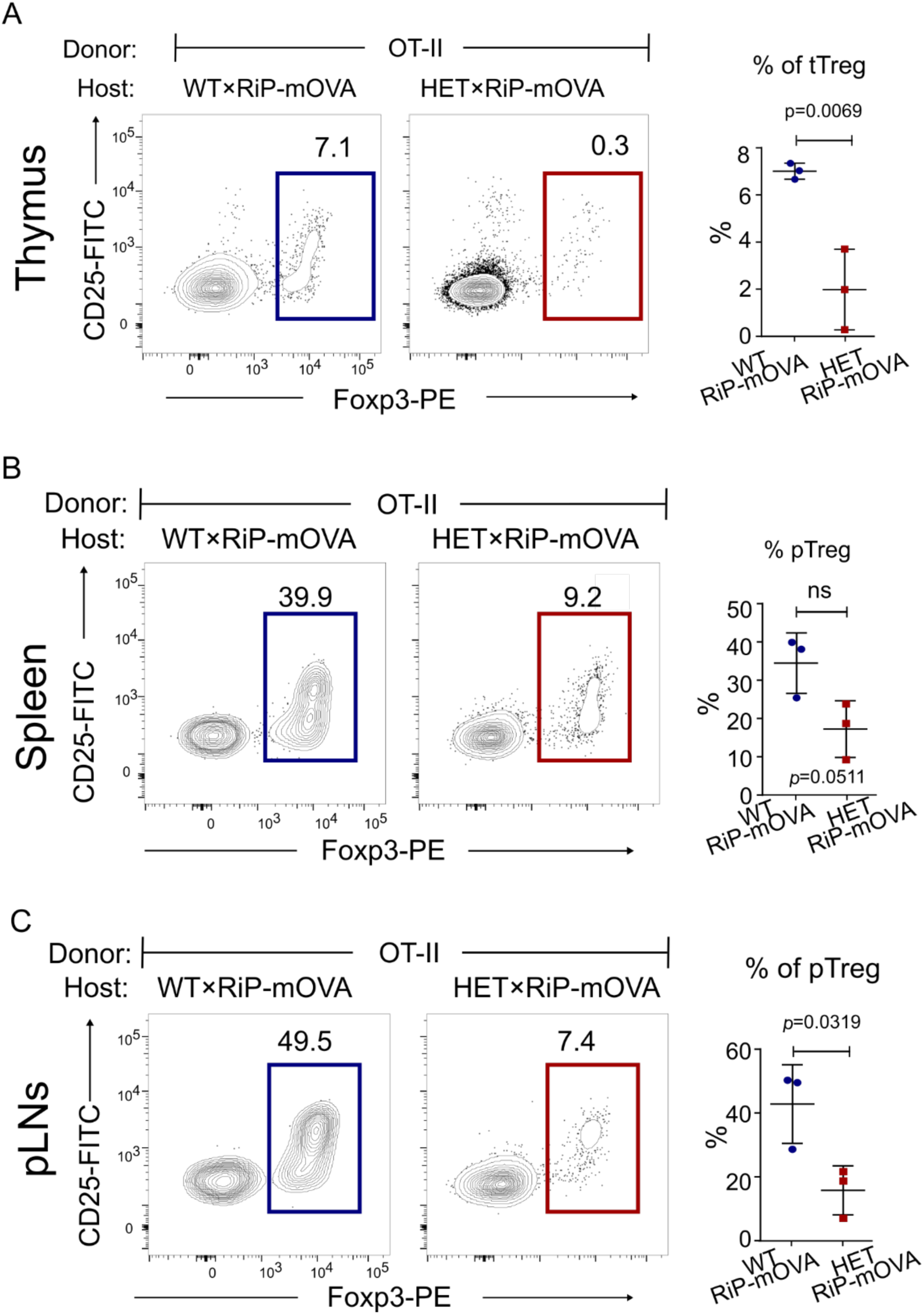
Mutant Copa impairs the generation of antigen-specific Tregs. A. left: Intracellular Foxp3 within CD4SP thymocytes. right: Percentages of Foxp3^+^ CD4SP among total CD4SP (WT×RiP-mOVA, *n* = 3; HET×RiP-mOVA, *n* = 3). B. left: Representative flow plot of intracellular Foxp3 levels within the reconstituted splenic CD4^+^ T cells. right: Percentages of pTregs among the reconstituted splenic CD4^+^ T cells (WT×RiP-mOVA, *n* = 3; HET×RiP-mOVA, *n* = 3). C. left: Intracellular Foxp3 within the reconstituted CD4^+^ T cells from pancreatic lymph nodes. right: Percentages of Foxp3^+^ CD4^+^ cells among reconstituted CD4^+^ T cells in pancreatic lymph nodes (WT×RiP-mOVA, *n* = 3; HET×RiP-mOVA, *n* = 3). Data are mean ± *SD*. Unpaired, parametric, two-tailed Student’s *t*-test was used for statistical analysis. *p* < 0.05 is considered statistically significant. ns: not significant.

### T cells from Copa^E241K/+^ mice cause ILD

Following our discovery that *Copa^E241K/+^* mice possess a defect in thymic tolerance, we hypothesized that autoreactive T cells may be pathogenic drivers of lung disease in our model. In support of this, we observed large numbers of tissue infiltrating CD4^+^ T cells in lung tissue sections harvested from *Copa^E241K/+^* mice (fig. S2D) and COPA syndrome subjects (*1*). To directly test whether T cells from *Copa^E241K/+^* mice mediate lung disease, we magnetically sorted T cells harvested from lymph nodes and spleens of WT and *Copa^E241K/+^* mice and adoptively transferred them into immunodeficient animals. Lungs from recipient mice demonstrated significant peribronchiolar and perivascular mononuclear infiltrates, remarkably similar to the spontaneous lung disease in unmanipulated *Copa^E241K/+^* mice (Fig. 8A-C). Taken together, these findings support a model whereby a defect in central tolerance caused by mutant Copa expression in the thymus leads to the escape of lung autoreactive T cells into the bloodstream that travel to the lung and cause ILD.

**Figure 8.**
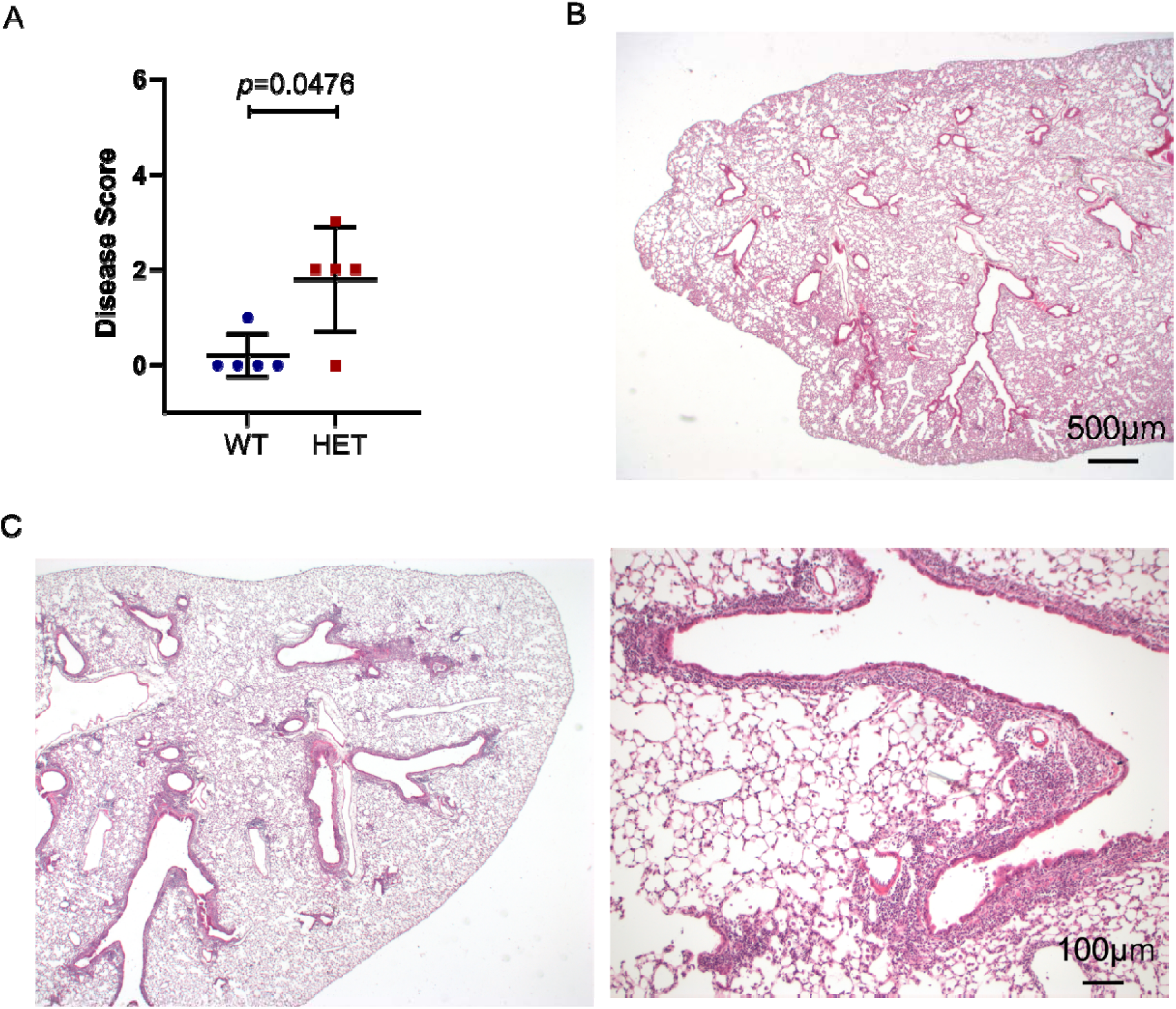
T cells from *Copa^E241K/+^* mice cause ILD. A. Disease scores of lung sections from recipient mice after adoptive transfer of T cells from WT or HET animals (WT transfer, *n* = 5; HET transfer, *n* = 5). B. Representative low power image of recipient mouse lung after WT T cell transfer. C. Recipient mouse lung after HET T cell transfer at low power (left) and high power (right) demonstrates peribronchial infiltration. Mann–Whitney U test was used in A. *p* < 0.05 is considered statistically significant. Low power images taken at 2x, scale bar = 500 μM. High power image taken at 10x, scale bar = 100 μM.

## Discussion

Here we describe a novel knock-in mouse modeled on one of the *COPA* missense mutations identified in patients. We found that mutant mice spontaneously developed pulmonary disease that closely mirrored the lung pathology found in patients. Mice also exhibited significant elevations in activated effector memory T cells, cytokine-secreting T cells and single positive thymocytes. Using bone marrow chimera and thymic transplant experiments we carefully mapped these abnormalities to mutant Copa in the thymic epithelium. Importantly, we discovered that a key mechanism for the breakdown of immune tolerance in our model was that a defect in thymic T cell selection brought forth two highly consequential outcomes: the escape of autoreactive T cells into peripheral lymphoid organs combined with a reduction in antigen-specific Tregs that are important for controlling inflammation. Finally, T cells from *Copa^E241K/+^* mice were shown to be pathogenic through adoptive transfer experiments that demonstrated their capacity to mediate lung disease. Given that ILD is the clinical characteristic that imparts the greatest impact on treatment and prognosis in COPA syndrome (*2*), *Copa^E241K/+^* mice should serve as an important model for studying how T cells cause the disease and how they might be targeted for successful treatment of patients.

To date, the underlying mechanisms of COPA syndrome have remained elusive. Although described as an autoimmune disorder, patients also have features of autoinflammation (*1, 21*) and it has been unclear whether *COPA* mutations result in dysfunction of adaptive immunity, innate immunity or both (*22*). Our study provides strong evidence that a key step in the initiation of disease in COPA syndrome is a breakdown in central tolerance in the thymus. This has important implications for guiding patient therapy because treatments for autoinflammation typically target IL-1 and TNF cytokine axes (*23*) whereas those for autoimmunity focus on dampening adaptive immune cell responses by B and T cells (*24*). Our observation that T cells appear to be a primary driver of lung disease suggests that therapies targeting T cells might be effective at treating ILD in COPA syndrome. Indeed, a recent study of COPA syndrome observed that use of purine synthesis inhibitors such as mycophenolate mofetil (MMF) that inhibit activated T lymphocytes (*25*) has shown some success at controlling pulmonary inflammation (*2*). Further work to understand which CD4 effector T cell populations are most responsible for disease in *Copa^E241K/+^* mice may provide new avenues for other T cell targeted therapies (*26, 27*).

Our study provides deeper insight into the pathogenesis of autoimmune-associated ILD and reinforces the link between ILD and defective thymic tolerance (*28*). A recent study found that patients with impaired central tolerance due to deficiency in the transcriptional regulator AIRE had a much higher prevalence of ILD than previously reported (*29*). Investigations of Aire-deficient patients and mice were used to define the antigen specificity of autoimmune-mediated ILD and identify autoantibody biomarkers to detect patients with autoimmune ILD (*11*). Similar studies should be undertaken in our model to determine if mutant Copa leads to impaired presentation of lung antigens in the thymus and eventually tissue-specific autoantibodies that mark the presence of ILD in COPA syndrome. Because Copa signifies a new molecular link between impaired central tolerance and ILD that is distinct from and independent of Aire function, this raises the possibility that the COPA model can be used to define new lung autoantigens targeted in ILD pathogenesis.

Although our study shows that adoptive transfer of T cells from *Copa^E241K/+^* mice were sufficient to cause ILD, the observation that some patients receiving maintenance immunosuppression experience unrelenting progression of lung fibrosis (*2*) suggests there may be a lung-intrinsic role to the disease. Importantly, studies in rheumatoid arthritis and scleroderma suggest that dysfunction within cells of the lung stroma contribute to ILD in these disorders (*30, 31*). As a result, antifibrotic medications currently approved for idiopathic pulmonary fibrosis are being studied in clinical trials as treatment for ILD associated with scleroderma and RA (*32, 33*). Similar investigations should be undertaken to determine whether mutant Copa exerts an effect within the lung parenchyma to better understand whether combination therapy with mycophenolic acid and antifibrotic drugs such as nintedanib might be beneficial in COPA syndrome, similar to what is being trialed in scleroderma ILD (*32*).

Remarkably, *Copa^E241K/+^* mice spontaneously recapitulated the lung disease and T cell activation observed in patients. The lack of other significant autoimmune features including autoantibodies and inflammatory joint disease may be a consequence of the mouse microbiome or because our mice our housed under specific pathogen free conditions, two variables that have been shown in several studies to impact the incidence and severity of clinical disease (*34, 35*). It is also well-established that autoimmunity within murine models is affected by mouse strain, likely because of differences in MHC haplotype (*36*). Although *Copa^E241K/+^* mice developed several immunologic perturbations shortly after birth, the lung disease appeared primarily in aged animals. We speculate that lung disease in human COPA syndrome may be triggered by viral infections that can vary from patient to patient. This may account for the variability in ILD severity and incomplete penetrance of disease in unaffected carriers of *COPA* mutations. It will be interesting to test this hypothesis directly in *Copa^E241K/+^* mice using viral infections or other experimental lung injury models to learn the impact of these challenges on ILD and systemic autoimmunity. It remains possible that *Copa^E241K/+^* mice on other background strains are more susceptible to developing earlier and more severe clinically-apparent disease including autoantibodies (*36*).

Our study raises several questions that should be explored in future investigations. First, the cellular mechanisms by which mutant Copa in thymic epithelial cells leads to a defect in negative selection remains unclear. Disruptions in COPI trafficking have been shown to alter endolysosomal function and macroautophagy (*1, 37–39*), cellular processes important in the proteolysis and loading of self-proteins onto MHC molecules (*40, 41*). We previously showed that cell lines derived from COPA syndrome patients demonstrate a constellation of morphological and functional findings that suggest impaired macroautophagy (*1*). Given the importance of macroautophagy to thymocyte selection and tolerance (*13*), it is possible that disruption of macroautophagy function in the thymus by mutant Copa may underlie the defect in negative selection we observed. Second, although our study confirms that the thymic epithelium is defective in COPA syndrome, this does not preclude other cell types from contributing to the disorder. Future investigations should evaluate whether disturbances in Copa function within other immune cells or tissues have an important role in disease pathogenesis.

Our findings have several important implications for patients. Given the significance of T cell-mediated lung injury, a range of medications targeting T cell responses should be studied to determine which are most effective. One of the major downsides to many of the immunosuppressive drugs currently in use, however, is that they globally suppress the immune system and place patients at increased risk of developing opportunistic infection and malignancy (*26*). Future investigations should evaluate whether antigen-specific tolerogenic strategies with the potential to exert a more potent effect on the disease can be used for treatment, particularly since these would avoid some of the more damaging side effects of global immune suppression. For instance, given the marked loss of antigen-specific Tregs we observed in *Copa^E241K/+^* mice, regulatory T cell based therapy may be an exciting area for future research once the critical autoantigens targeted in disease are defined (*27, 42*), especially in light of recent evidence which shows an important role for Tregs in constraining lung fibrosis (*43*).

Because mutant Copa in the thymic stroma plays a central role in the generation of pathogenic T cells, developing methods to repair thymic epithelial cells directly might in theory be highly effective. Although significant barriers remain, one can envision a day when evolving technologies such as CRISPR/Cas9 gene-editing may be used to correct a *COPA* mutation in the thymus (*44*). Work is currently underway to study how CRISPR/Cas9 tools combined with induced pluripotent stem (iPS) cells can be employed to generate a gene-edited thymus. In principle, iPS cells from COPA syndrome patients could be edited with CRISPR/Cas9 tools to correct the disease-causing mutation and then differentiated into functional thymus cells. Once achieved, the iPS-derived thymus cells could be transplanted back into the patient in an attempt reestablish normal thymic selection and development (*45*).

In conclusion, we established a new mouse model of COPA syndrome and identified Copa as a novel molecular mechanism regulating tolerance in the thymus. We demonstrate that mutant Copa specifically within thymic epithelial results in a striking defect in thymocyte selection that results in the release of autoreactive T cells into peripheral organs and the loss of antigen-specific Tregs. Taken together, our data provides compelling evidence that a key step in the initiation of autoimmunity in COPA syndrome patients is a breakdown in central tolerance of the thymus.

## Materials and Methods

### Copa^E241K^ targeting construct and mice strains

The BAC RP24-64H24 (Children’s Hospital Oakland Research Institute) containing mouse *Copa* of C57Bl/6 origin served as template. To generate the 5’ homology arm, the locus containing exons 9 and 10 was amplified with NotI and PacI flanking the amplicon and then TOPO cloned into pCR-BluntII-TOPO (Life Technologies). The codon encoding residue 241 of Copa was mutated from glutamic acid to lysine via Quikchange Lightning site directed mutagenesis (Agilent). Following NotI and PacI digestion, the 5’ homology arm was ligated into pEasyFloxDTA (gifted by Dr. Mark Anderson, UCSF). The locus containing exons 11 through 15 of *Copa* was amplified off RP24-64H24 with FseI flanking ends and ligated into pEasyFloxDTA with Infusion recombination (Takara Bio USA) to form the 3’ homology arm.

Twenty micrograms of linearized vector were electroporated twice into 20 million JM8A3.N1 embryonic stem cells (UCSF ES Cell Targeting Core). The ES cells were chosen for C57Bl/6 origin and agouti coat color mutation. ES cells were positively selected with neomycin and negatively selected with diphtheria toxin A, resulting in 252 clones.

After determining successful recombination with PCR analysis, one positive clone was expanded and screened with Southern blotting. Genomic DNA was digested with BglII and KpnI, size separated with agarose gel electrophoresis, and transferred to Amersham Hybond N+ (GE Healthcare Life Sciences). Probe radiolabeled with α-^32^P dATP (Perkin Elmer) was hybridized to the membrane in QuikHyb (Agilent) and followed by exposure to X-ray film.

The expanded clone was injected into C57Bl/6 blastocysts and transplanted into pseudo pregnant females (Gladstone Transgenic Gene Targeting Core). Fifteen founders were born. Tissue biopsy was collected from chimeric founders to genotype by Sanger sequencing. Following germline transmission of *Copa^E241K^*, mice were crossed to B6.Cg-Tg(Sox2-cre)1Amc/J mice (Jackson Laboratory) for Cre recombinase mediated removal of the *Neo* cassette. Resulting mice were crossed to C57Bl/6J (Jackson Labs) for two generations to expand the colony.

B6.Cg-Tg(TcraTcrb)425Cbn/J (OT-II), C57BL/6-Tg(TcraTcrb)1100Mjb/J (OT-I), C57BL/6-Tg(Ins2-TFRC/OVA)296Wehi/WehiJ (RIP-mOVA), B6.SJL-*Ptprc^a^ Pepc^b^*/BoyJ (B6 Cd45.1), and B6.Cg-*Foxn1^nu^*/J (B6 Nude) mice were purchased from the Jackson Laboratory. B6;D2-TCR LCMV (P14) mice were provided by Dr. Mehrdad Matloubian at UCSF. All mice were maintained in the specific pathogen free (SPF) facility at UCSF, and all protocols were approved by UCSF’s Institutional Animal Care and Use Committee (IACUC).

#### Flow cytometry and antibodies

Single cell suspension of thymocytes and splenocytes were prepared by mechanically disrupting the thymus and spleen. Cells were filtered through 40 µm filters (Falcon) into 15 ml conical tubes and maintained in RPMI 1640 containing 5% FBS on ice. Splenocytes were further subjected to red blood cells lysis (Biolegend). One aliquot of each single cell suspension was mixed with AccuCheck Counting Beads (Invitrogen, PCB100) and analyzed by flow cytometry. The total cell number was back calculated according to manufacturer’s protocol. For evaluation of surface receptors, cells were blocked with 10 µg/ml anti-CD16/32 for 15 minutes at room temperature and then stained with indicated antibodies in FACS buffer (PBS, 2%FBS) on ice for 40 minutes.

For intracellular cytokine detection, freshly isolated splenocytes were stimulated by PMA (Sigma) and ionomycin (Sigma) at 37**°**C in the presence of brefeldin A (Biolegend). Four hours following stimulation, cells were collected and stained with Ghost Dye Violet 450 (Tonbo) followed by surface staining. Cells were then washed, fixed at room temperature for 15 minutes (PBS, 1% neutral buffered formalin), permeabilized and stained with antibodies against cytokines on ice (PBS, 2% FBS, 0.2% saponin). For nuclear protein detection, cells were stained with the eBioscience Foxp3 staining kit following the manufacturer’s protocol. All samples were acquired on LSRFortessa or FACSVerse (BD Bioscience) and analyzed using FlowJo software V10.

Antibodies to the following were purchased from Biolegend: CD8 (53-6.7), CD3 (145-2C11), CD62L (MEL-14), IFNγ (XMG1.2), CD45 (30-F11), CD80 (16-10A1), B220 (RA3-6B2), Ly-51 (6C3), Epcam (G8.8), TCR Vβ8.1,8.2 (KJ16-133.18), TCR Vβ5.1,5.2 (MR9-4), TCR Vα2 (B20.1); from eBioscience: CD45.1 (A20), IL17A (eBio17B7), IL13 (eBio13A), Foxp3 (FJK-16S), Nur77 (12.14), MHCII (I-A) (NIMR-4); from Tonbo: CD69 (H1.2F3), B220 (RA3-6B2), CD4 (GK1.5), CD25 (PC61.5), MHCII (I-A/I-E) (M5/114.15.2), TCRβ (H57-597), CD44 (IM7), CD45.2 (104); and from Thermo Fisher Scientific: CD8 (5H10). The CD16/CD32 (2.4G2) antibody was obtained from UCSF’s Hybridoma Core Facility.

#### ELISA

Total IgG level were measured using mouse IgG ELISA kit (Bethyl Laboratories Inc, E90-131). 96 well flat-bottomed plates (Nunc, #439454) were coated with anti-IgG-FC overnight at room temperature. Plates were washed 5 times with washing buffer (PBS, 0.05% Tween), and then blocked with blocking buffer (PBS, 2%FBS, 0.05% Tween). Mice serum were diluted with blocking buffer at 1:10000, and added into each well for a 2h incubation at room temperature. After 5 washes, HRP-anti mouse IgG (1:3000 in blocking buffer) were added into each well for a 1-hour incubation. Plates were washed for 5 times, developed using TMB solution (R&D Systems, #DY999) and read at 450nm.

Anti-dsDNA ELISA were performed as previously described (*46*). 96 well flat-bottomed plates (Falcon, #351172) were coated with linearized dsDNA (HincII digested pUC19 plasmid) overnight at room temperature. Plates were washed and blocked. Mice serum were diluted with blocking buffer at 1:160, and added into each well for a 2-hour incubation at room temperature. After 5 washes, HRP-anti mouse IgG (1:3000 in blocking buffer) were added into each well for a 1-hour incubation. Plates were washed and developed using TMB solution and read at 450nm.

For anti-nRNP ELISA, 96 well flat-bottomed plates (Nunc, #439454) were coated with Sm/RNP (ImmunoVision, #SRC-3000) overnight. Plates were washed and blocked. Mice serum was diluted with blocking buffer at 1:40, and added into each well for a 2-hour incubation at room temperature. After 5 washes, HRP-anti mouse IgG (1:3000 in blocking buffer) were added into each well for a 1-hour incubation. Plates were washed and developed using TMB solution and read at 450nm.

#### Bone marrow chimeras

To deplete bone-marrow in host animals, 3-4-month-old mice were given 2 doses of γ-irradiation (550 rads each time) with a 12-hour interval. Bone marrow cells in femurs and tibias were harvested from 8-week-old, gender-matched donor mice. T cells were depleted using anti-CD90.2 antibody (Biolegend, 30-H112) and Dynabeads (Invitrogen, 65601). Ten million bone marrow cells were suspended in 200 µl PBS and intravenously introduced into the hosts. Chimeric mice were maintained under SPF conditions and analyzed 6-8 weeks after the reconstitution.

#### Thymic transplantation

Fetal thymi were isolated from E15.5 embryos and seeded into fetal thymus organ culture (FTOC) (*10*) 1.35 mM 2-deoxyguanosine (Sigma) was added to the culture medium to deplete thymocytes. 7 days later, the empty thymic lobes were collected and washed with sterile PBS for 4 hours and then transplanted under the kidney capsule of gender matched Nude mice. The reconstituted thymi were harvested 6-8 weeks after transplantation, and thymocytes were analyzed by flow cytometry.

#### Isolation and transfer of CD3^+^ T cells

Single cell suspension from lymph nodes and spleens were prepared as described above. CD3^+^ T cells were isolated using MojoSort negative selection kit (Biolegend, #480031). Derived cells were counted and purity of the cells were checked by flow cytometry. 10 million CD3^+^ T cells were suspended in 150µl PBS and introduced into *Rag2^-/-^* mice through i.p. injection. Three months after the adoptive transfer, lung tissues were harvested from the *Rag2^-/-^* recipients and subjected to histological analysis.

#### Quantitative RT-PCR

Thymic epithelial cells were prepared as described previously (*47*) and then sorted using a FACS Aria II (BD Biosciences). RNA was isolated using Dynabeads® mRNA Purification Kit (Invitrogen) and reverse transcribed using SuperScript III Reverse Transcriptase and oligo d(T)_16_ primers (Invitrogen). qPCR was performed with TaqMan Gene Expression assays from Applied Biosystems (*Gapdh*, Mm99999915_g1; *Aire*, Mm00477461_m1). A Custom TaqMan assay (*17*) was used for *mOVA* transcripts.

For *Copa* transcript detection, RNA was prepared from the skin of indicated mice using RNeasy Mini kit (Qiagen). SuperScript III Reverse Transcriptase and oligo d(T)_16_ primers (Invitrogen) were used to synthesize cDNA. qPCR was performed with TaqMan Gene Expression assays from Applied Biosystems (*Copa*, Mm00550231_m1; *Actb*, 4352933E).

A Custom TaqMan assay was used for specific Copa transcripts: fwd: 5’-CAAGTGAAGATCTGGCGTATGAATG-3’, rev: 5’-AAAACAGCACAAGACACATTGTTGT-3’, probe for wildtype Copa allele: 5’-Vic-AGGCATGGGAGGTTGA-MGB-3’, probe for mutant Copa allele: 5’-Fam-AGGCATGGAAGGTTGA-MGB-3’.

Data were acquired with Bio-Rad CFX96 Real Time Machine.

#### Histology

Lung tissue was harvested and fixed in 4%PFA overnight, followed by fixation in 70% Ethanol for 24 hours before embedding, sectioning and H&E staining. The following grading system was used to score the infiltration: grade 0, no infiltrates; grade 1, few perivascular and peribronchiolar mononuclear infiltrates; grade 2, frequent perivascular and peribronchiolar mononuclear infiltrates; grade 3, diffuse or dense perivascular and peribronchiolar mononuclear infiltrates; grade 4, diffuse perivascular and peribronchiolar mononuclear infiltrates, interstitial pneumonia and architectural distortion. For immune cell subtyping, fresh lung tissues were inflated with a mixture of optimal cutting temperature compound and sucrose, and were snap frozen on dry ice. 8µm sections from the frozen lungs were cut on a cryostat. Antibodies specific for CD4, CD8 and B220 (BD Bioscience) and a DAB Kit (Vector Laboratories) were used for the immunohistochemistry detection.

#### Western Blot

Protein lysates were subjected to SDS-PAGE and transferred to PVDF membranes (Millipore). Membranes were blocked and probed with one of the following antibodies: a rabbit antibody to COPA (Sigma, HPA028024) or a mouse antibody to GAPDH (Santa Cruz Biotechnology, sc-32233). Bands were detected by incubation with secondary HRP-coupled antibodies (Jackson Immunoresearch) and SuperSignal West Femto (Pierce) chemiluminescence.

#### Statistical analysis

All statistical analysis was performed using Prism 7 (GraphPad Software). Mann–Whitney *U* test was used to evaluate statistical significance of the lung disease scores. Unpaired, parametric, two-tailed Student’s *t*-test was used to evaluate the statistical significance between two groups. All graphs display mean ± standard deviation (*SD*), and *p* < 0.05 was considered statistically significant.

## Supplementary Materials

Fig. S1. Schematic for generation of *Copa^E241K/+^* mice and validation of gene targeting.

Fig. S2. Copa^E241K/+^ mice develop lymphocytic infiltration of lung but no joint disease.

Fig. S3. Copa^E241K/+^ mice have normal percentages of splenic B and T cells and no autoantibodies.

Fig. S4. Mutant Copa in thymic stroma causes an increase in SP thymocytes despite apparent normal positive selection in Copa^E241K/+^ mice

## Author contributions

Z.D. designed and performed experiments, analyzed data and wrote the manuscript. C.S.L. performed experiments and analyzed data. F.O.H. and K.M.W performed experiments. J.S. and K.D.J. analyzed data. A.K.S. directed the study and wrote the manuscript.

## Data and materials availability

All data is available in the main text or the supplementary materials.

