## Supplemental data for "A defect in thymic tolerance causes T cell-mediated autoimmunity in a murine model of COPA syndrome"

**Supplementary Materials**

#### Supplementary Figure 1

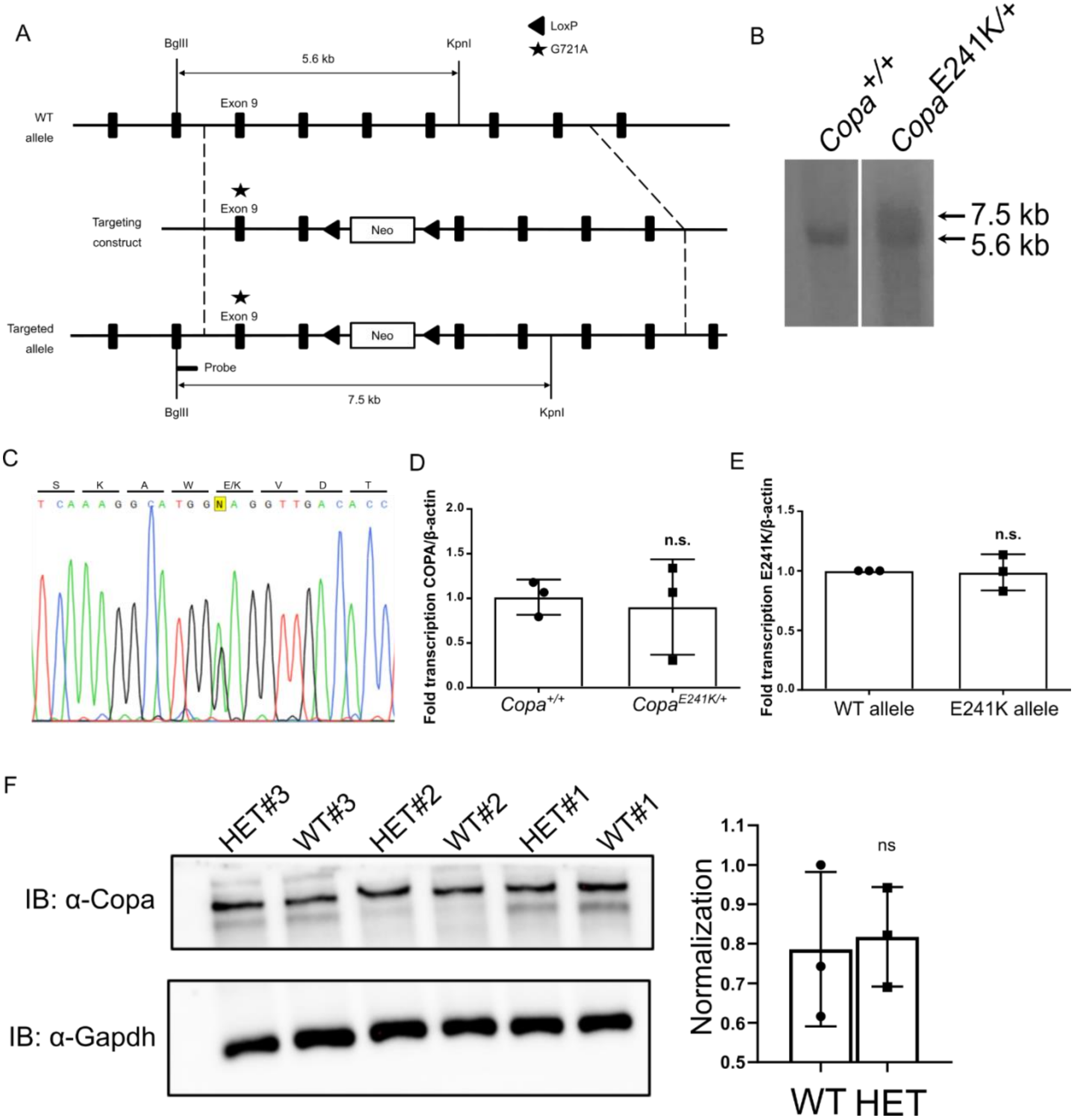

**Fig. S1: Schematic for generation of *Copa*$_{E241K/+}$ mice and validation of gene targeting.**

A. Schematic of the targeting vector used to knock-in the E241K mutation into exon 9 of the *Copa* gene. The Neomycin cassette is flanked by LoxP sites. The dashed lines indicate the ends of the targeting construct's homology.

B. Southern blot of genomic DNA from ES cells targeted with the E241K/G721A knock-in construct and digested with BglII and KpnI.

C. The presence of the mutation was confirmed by Sanger sequencing in ES cells that were positive by Southern blot.

D. Real time PCR measurement of total *Copa* mRNA level in WT and *Copa*$_{E421K/+}$ mice (WT, $n$ = 3; HET, $n$ = 3).

E. mRNA level of wild type *Copa* and *Copa*$_{E241K}$ alleles in heterozygous *Copa*$_{E421K/+}$ mice ($n$ = 3).

F. left: Western blot showing the Copa protein level in the thymocytes from indicated mice (WT, $n$ = 3; HET, $n$ = 3). right: Quantification of the western blot.

Data are mean ± *SD*. Unpaired, parametric, two-tailed Student's *t*-test was used for statistical analysis. $p < 0.05$ is considered statistically significant. ns: not significant.

### Supplementary Figure 2

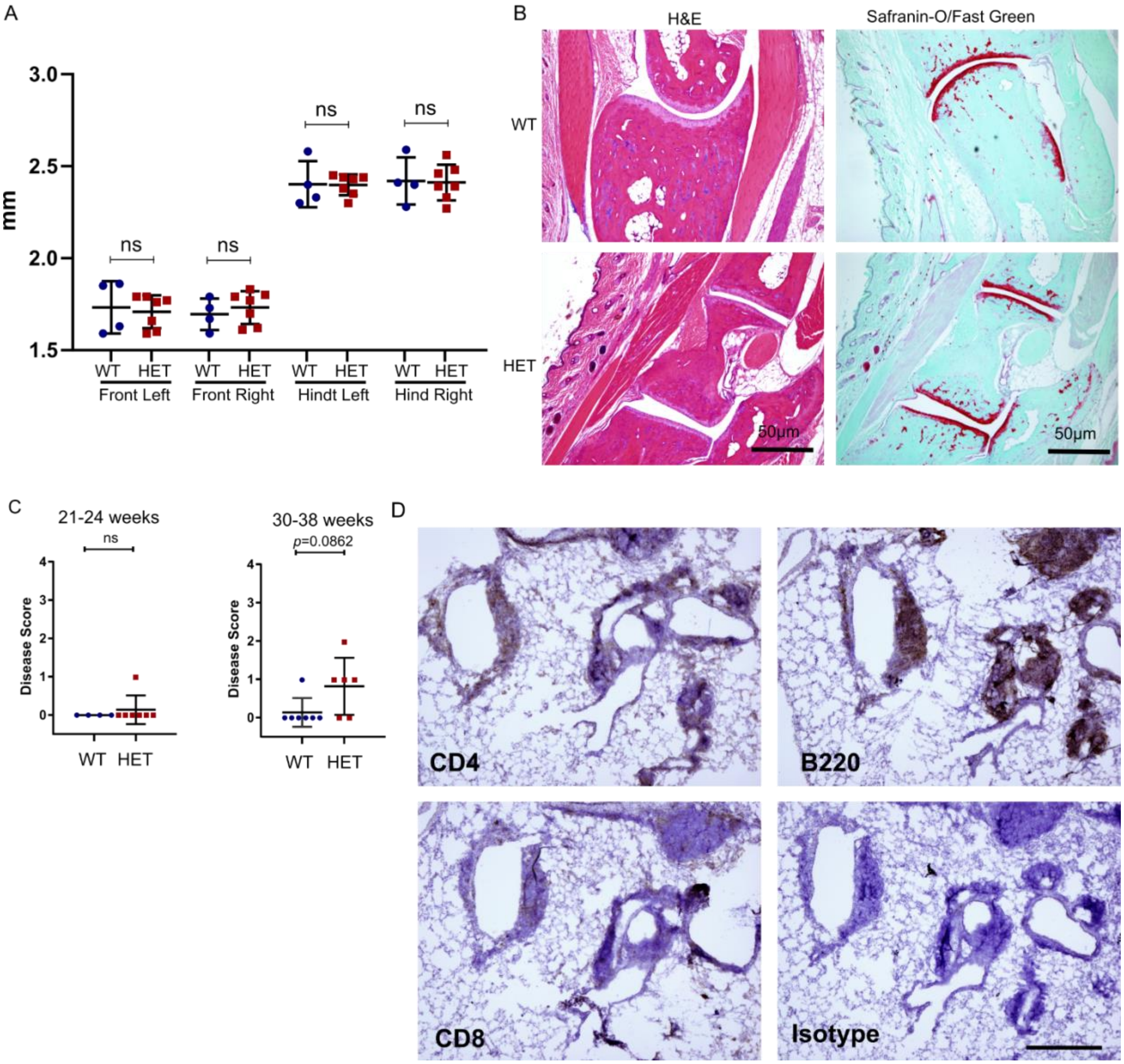


**Fig. S2: *Copa*$^{E241K/+}$ mice develop lymphocytic infiltration of lung but no joint disease**

A. Ankle thickness of the wild type and *Copa*$^{E241K/+}$ mice (6-month-old litter mates: WT, $n$ = 4; HET, $n$ = 7).

B. left: H&E staining of the ankle sections of the hind legs. right: Safranin-O/Fast green staining of the ankle sections of the hind legs.

C. Disease scores of lung sections from the 5-6-month-old (Littermates: WT, *n* = 4; HET, *n* = 7) and 7-9-month-old (Littermates: WT, *n* = 7; HET, *n* = 6) mice.

D. Representative image of the IHC staining of the lung section from *Copa*$^{E241K/+}$ mice.

Data are mean ± *SD*. Unpaired, parametric, two-tailed Student's *t*-test was used for statistical analysis in A. $p < 0.05$ is considered statistically significant. ns: not significant. B and D taken at 4x magnification, scale bar = 50μm.

#### Supplementary Figure 3

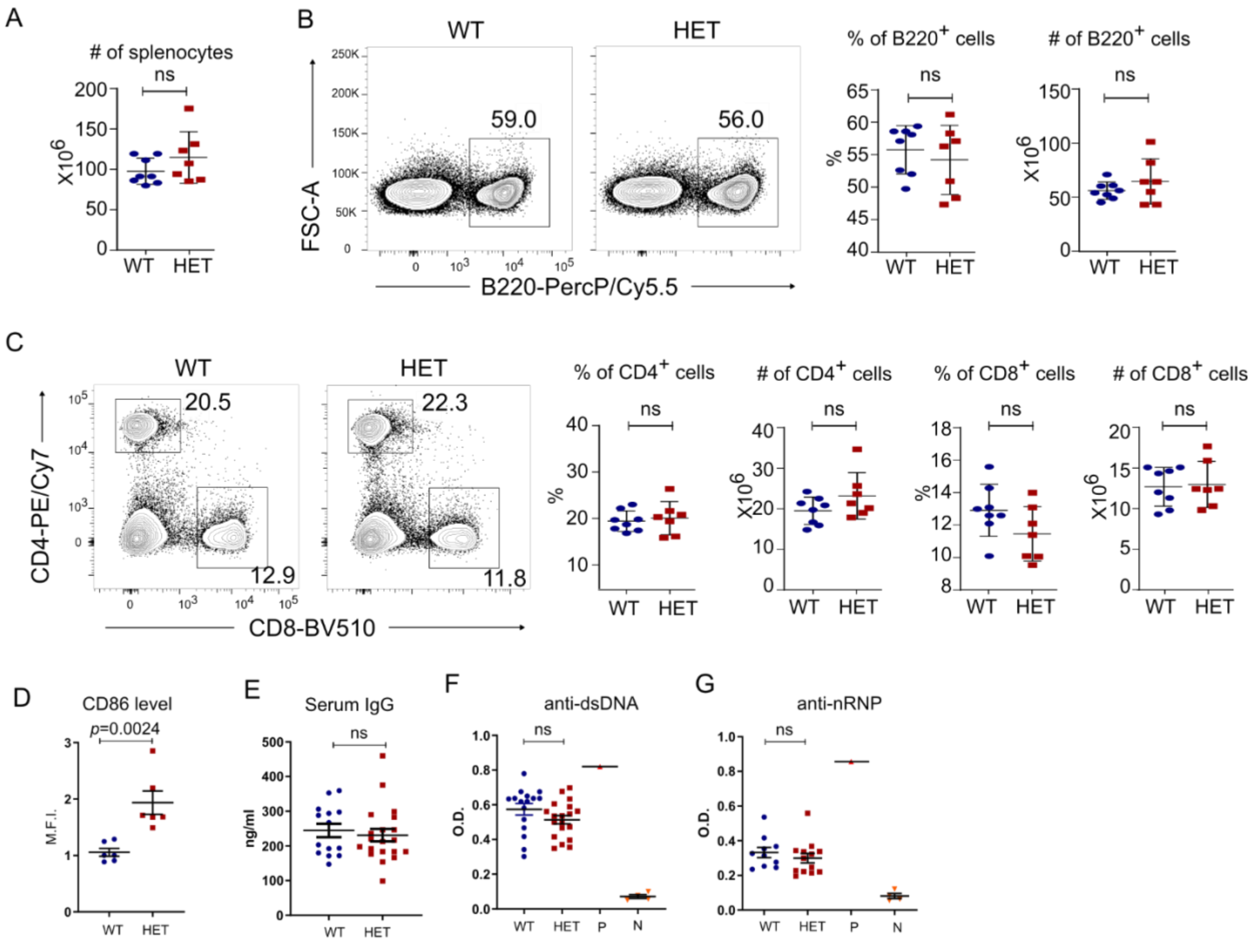

**Fig. S3: *Copa*$^{E241K/+}$ mice have normal percentages of splenic B and T cells and no autoantibodies**

A. Cell counts of splenocytes (3-month-old littermates: WT, *n* = 8; HET, *n* = 7).

B. left: Representative flow plots showing percentage of B cells based on B220 expression. right: Percentage and cell counts of B220$^{+}$ cells among splenocytes (3-month-old littermates: WT, *n* = 8; HET, *n* = 7).

C. left: Representative flow plots showing the percentages of CD4$^{+}$ and CD8$^{+}$ T cells. right: Percentages and cell counts of CD4$^{+}$ and CD8$^{+}$ T cells among splenocytes (3-month-old littermates: WT, *n* = 8; HET, *n* = 7).

D. Quantification of the M.F.I. of CD86 on B cells. (3-month-old littermates: WT, *n* = 6; HET, *n* = 6)

E. Measurement of serum IgG level by Elisa (9-10-month-old littermates: WT, *n* = 14; HET, *n* = 20)

F. Serum anti-dsDNA IgG level by Elisa (9-10-month-old littermates: WT, *n* = 16; HET, *n* = 19). P: serum from a Lyn knock out mouse. N: PBS.

G. Serum anti-nRNP IgG level by Elisa (9-10-month-old littermates: WT, *n* = 10; HET, *n* = 13). P: serum from a Lyn knock out mouse. N: PBS.

Data are mean ± *SD*. Unpaired, parametric, two-tailed Student's *t*-test was used for statistical analysis. $p < 0.05$ is considered statistically significant. ns: not significant.

### Supplementary Figure 4

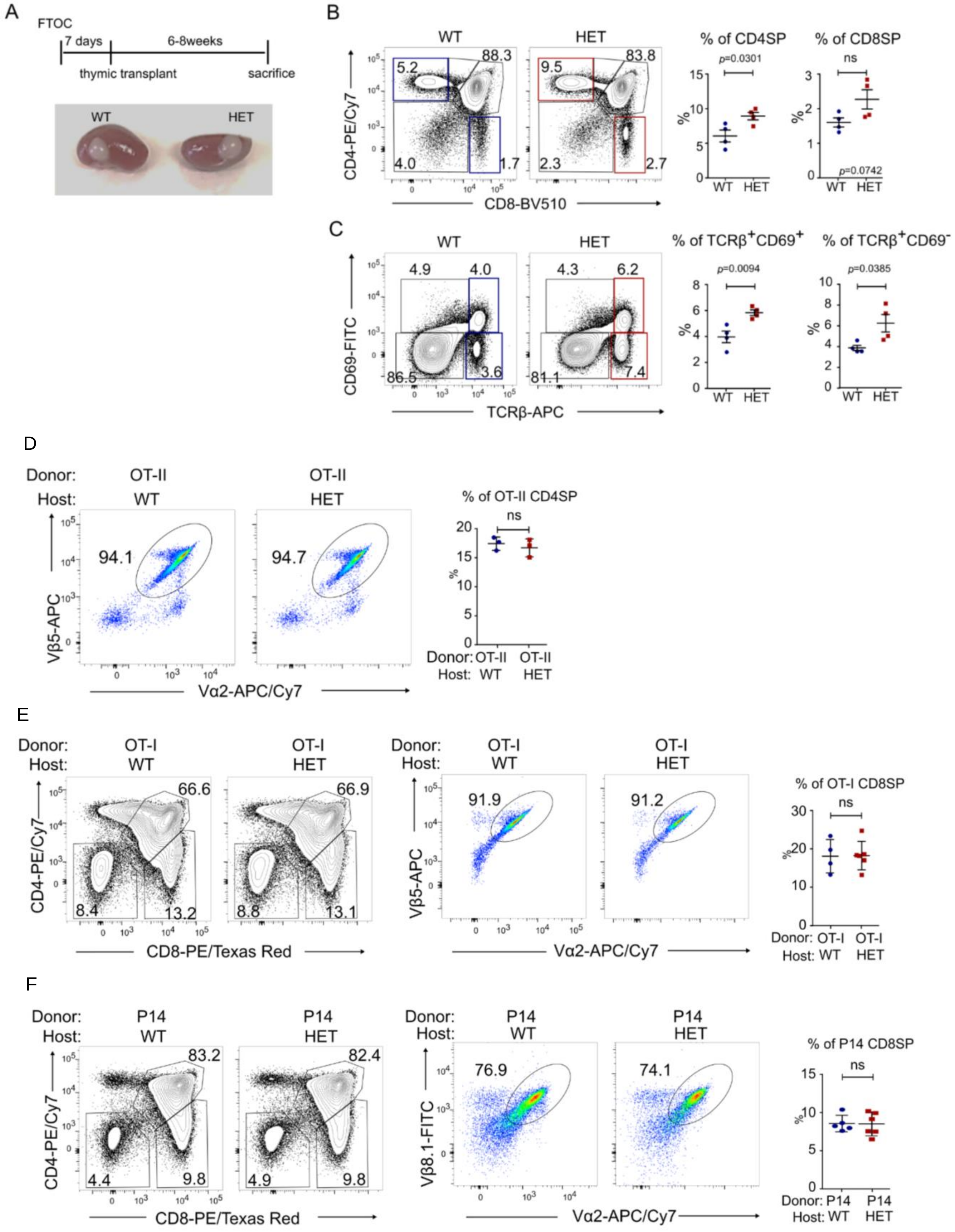

**Fig. S4. Mutant Copa in thymic stroma causes an increase in SP thymocytes despite apparent normal positive selection in *Copa*$^{E241K/+}$ mice**

A. top: Schedule from fetal thymus organ culture (FTOC) to transplantation to analysis. bottom: An image of grafted thymi.

B. left: CD4 and CD8 profile of thymocytes in transplanted thymi. right: Percentages of CD4SP and CD8SP (WT, *n* = 4; HET, *n* = 4).

C. left: CD69 and TCRβ profile of thymocytes in transplanted thymi. right: Percentages of TCRβ$^{+}$CD69$^{+}$ and TCRβ$^{+}$CD69$^{-}$ thymocytes (WT, *n* = 4; HET, *n* = 4).

D. left: Representative flow plots of Vα2 and Vβ5 on OT-II CD4SP thymocytes in WT and HET hosts. right: Percentages of OT-II CD4SP in the indicated hosts (WT, *n* = 3; HET, *n* = 3).

E. left: Representative CD4 versus CD8 flow plots of OT-I thymocytes in WT and HET hosts. middle: Flow cytometric measurement of Vα2 and Vβ5 on CD8SP thymocytes. right: Percentage of CD8SP in the indicated hosts (WT, *n* = 4; HET, *n* = 5).

F. left: Representative CD4 versus CD8 flow plots of P14 thymocytes in WT and HET hosts. middle: Flow cytometric measurement of Vα2 and Vβ8.1 on CD8SP thymocytes. right: Percentage of CD8SP in the indicated hosts (WT, *n* = 5; HET, *n* = 5).

Data are mean ± *SD*. Unpaired, parametric, two-tailed Student's *t*-test was used for statistical analysis. $p < 0.05$ is considered statistically significant. ns: not significant.
